# Stress-induced alterations of norepinephrine release in the bed nucleus of the stria terminalis of mice

**DOI:** 10.1101/335653

**Authors:** Karl T. Schmidt, Viren H. Makhijani, Kristen M. Boyt, Dipanwita Pati, Melanie M. Pina, Isabel M. Bravo, Jason L. Locke, Sara R. Jones, Joyce Besheer, Zoé A. McElligott

## Abstract

Stress can drive adaptive changes to maintain survival during threatening stimuli. Chronic stress exposure, however, may result in pathological adaptations. A key neurotransmitter involved in stress signaling is norepinephrine. Previous studies show that stress elevates norepinephrine levels in the bed nucleus of the stria terminalis (BNST), a critical node regulating anxiety and upstream of stress responses. Here, we use mice expressing channelrhodopsin in norepinephrine neurons to selectively activate terminals in the BNST, and measure norepinephrine release with fast-scan cyclic voltammetry. Mice exposed to a single restraint session show an identical norepinephrine release profile compared to that of unexposed mice. Mice experiencing five days of restraint stress, however, show elevated noradrenergic release across multiple stimulation parameters, and reduced sensitivity to the α_2_-adrenergic receptor antagonist idazoxan. These data are the first to examine norepinephrine release in the BNST to tonic and phasic stimulation frequencies, and confirm that repeated stress alters autoreceptor sensitivity.

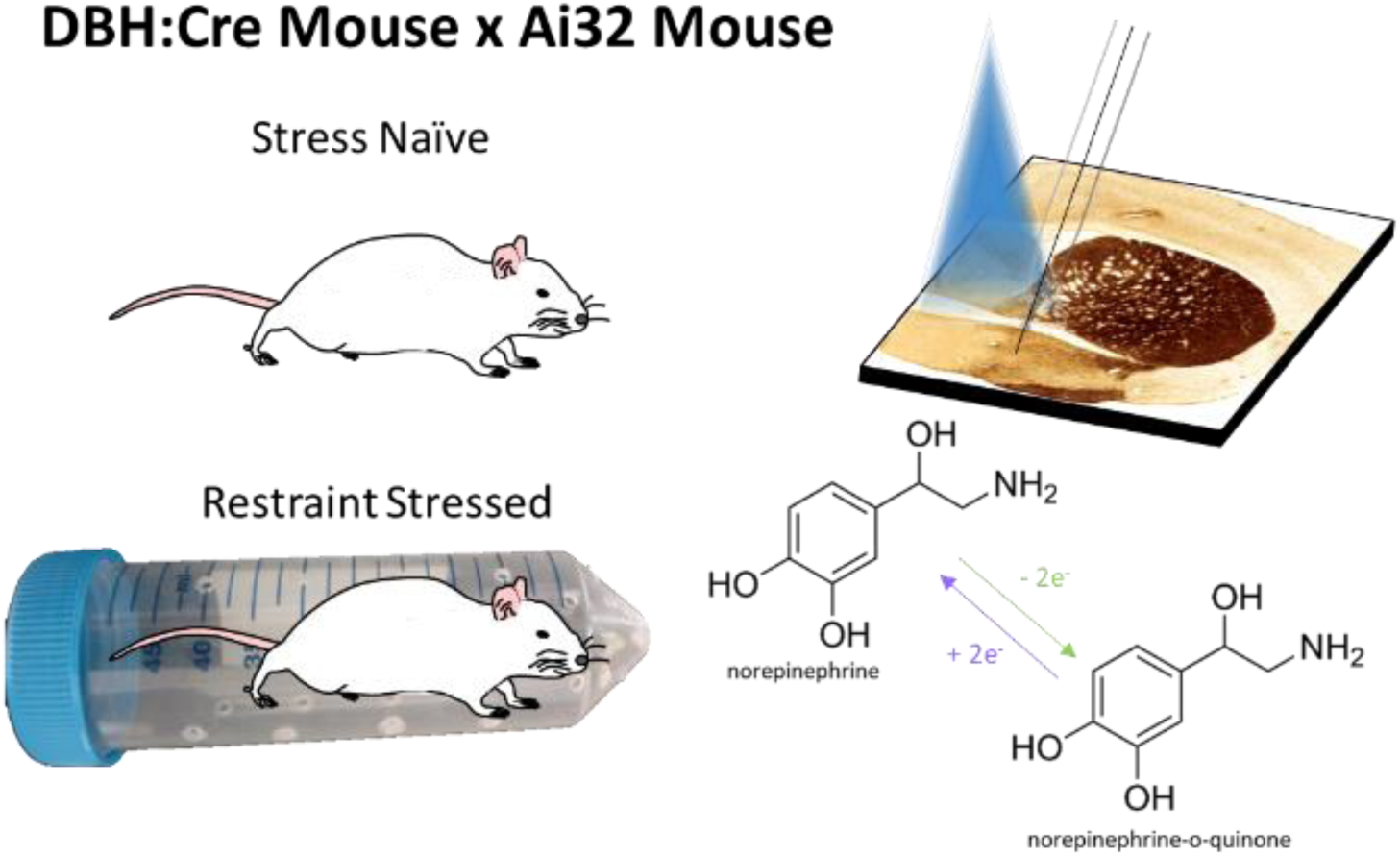

---

Stress is an important physiologic response to potentially harmful environmental stimuli. Under some conditions, however, such as chronic stress exposure or especially traumatic events, persistent alterations in the stress response occur, leading to abnormal and potentially maladaptive responses. Previous research has shown a central role of the catecholamine neurotransmitter norepinephrine (NE) in the physiological and behavioral stress responses^1^. NE release, in key downstream nuclei following stress has been implicated in a number of psychological disorders including addiction^2-3^, affective disorders^1^, and PTSD^1^.

Noradrenergic neurons project through two primary pathways in the brain: the dorsal noradrenergic bundle and ventral noradrenergic bundle. The pontine locus coeruleus (A6) gives rise to the dorsal noradrenergic bundle sending projections throughout the cortex, thalamus, hippocampus, cerebellum, and amygdala^5-7^. In contrast, the ventral noradrenergic bundle mainly arises from two nuclei, A1 of the rostral ventrolateral medulla and A2 of the nucleus of the solitary tract^6^, and projects mainly to hypothalamus, parabrachial nucleus, midbrain, and bed nucleus of the stria terminalis (BNST)^3, 6, 8^. Of particular interest is the projection from the medullary noradrenergic nuclei (i.e. A1/A2) to the BNST because it comprises the densest region of NE terminals in the brain^9-12^ and serves as a critical nucleus for processing affective state^13^. Furthermore, multiple stressors have been shown to increase NE release in the BNST, and blockade of α_1_-adrenergic receptors (α_1_-AR) in the BNST reduced circulating adrenocorticotropic hormone concentrations and anxiety-like behavior^14^. There appears to be a critical role of adrenergic receptors in the BNST in the response to stress, as chronic restraint stress prevents the ability of the region to express α1-AR dependent plasticity^15^ and multiple adrenergic receptors can drive release of corticotropin releasing factor within the BNST^15^ ^16^.

Previously, we and others have demonstrated that stress exposure modulates norepinephrine release in the BNST in rats^17-18^. Due to innervation of multiple biogenic amine pathways, and a reliance on electrical stimulation, these studies could not be performed at physiologically relevant frequency patterns^19-20^, and it was challenging to isolate catecholamine release from noradrenergic terminals^11^. Here we use a combination of optogenetics and fast-scan cyclic voltammetry (FSCV) to examine stress-induced alterations in NE release in the BNST. Using a mouse line expressing Cre-recombinase driven by the dopamine-beta-hydroxylase (DBH::Cre) promoter crossed with the Allen Institute’s “Ai32” mouse line^21^ that expresses channelrhodopsin (ChR2) in a Cre-recombinase dependent manner, we were able to selectively probe noradrenergic release in the BNST using optogenetically-driven stimulation with FSCV following various restraint stress exposure paradigms. We show that repeated restraint stress exposure, but not single stress exposure, enhances NE release in the BNST of mice. Our data suggests this phenomenon may be driven by alterations in α_2_-adrenergic receptor function.

## Results and Discussion

Before using the DBH:Cre(+/-)::Ai32(+/ +) mice to examine stress-induced neuroadaptations, we first characterized the noradrenergic system in this line. Transgene expression did not alter tissue content of NE in the BNST, cortex, striatum, hippocampus, or amygdala as measured by high-performance liquid chromatography (data not shown). Using *ex vivo* patch-clamp electrophysiology, we recorded the activity of ChR2-eYFP^+^ neurons of the A2 cell group in 300 μm coronal slices of the nucleus of the solitary tract. These neurons showed light-evoked action potentials at both low-tonic frequencies (10 pulses, 1 Hz, 5 ms, 473 nm; Fig. 1A) and high-phasic frequencies (15 Hz; Fig. 1B) with high fidelity (n=5 cells). Furthermore, we observed a dense fiber projection of noradrenergic neurons in the ventral BNST (Fig. 1C). Repeated optogenetic stimulations of equal parameters (20 pulses, 10 Hz, 5 ms, 473 nm) with 5 min between each stimulation induced stable NE release with an average peak concentration of 0.1689 μM NE at each stimulation (Fig. 1D). This concentration is in line with what was previously measured *in vivo* in rats with supraphysiological frequencies and higher pulse numbers of electrical stimulation at distal sites^11, 18, 22^. Previously reported dopamine release using a DAT::CRE line crossed with Ai32 mice indicated a decrease in optogenetically-evoked dopamine with repeated stimulations in the striatum^23^. As we do not see this consecutive decrease in our recordings, it may be a result of differences in Cre-driver line (as the DAT::Cre line has altered clearance mechanisms), brain region, or catecholamine. Furthermore, we observed stimulation dependent NE release with low concentrations (mean [NE] =0.037 μM; n = 8) detected following a single pulse of light and higher concentrations (mean [NE] = 0.243 μM; n = 6) following phasic burst-like (20 pulses, 15 Hz) stimulation (Fig. 1E, F, G, H).

**Figure 1.**
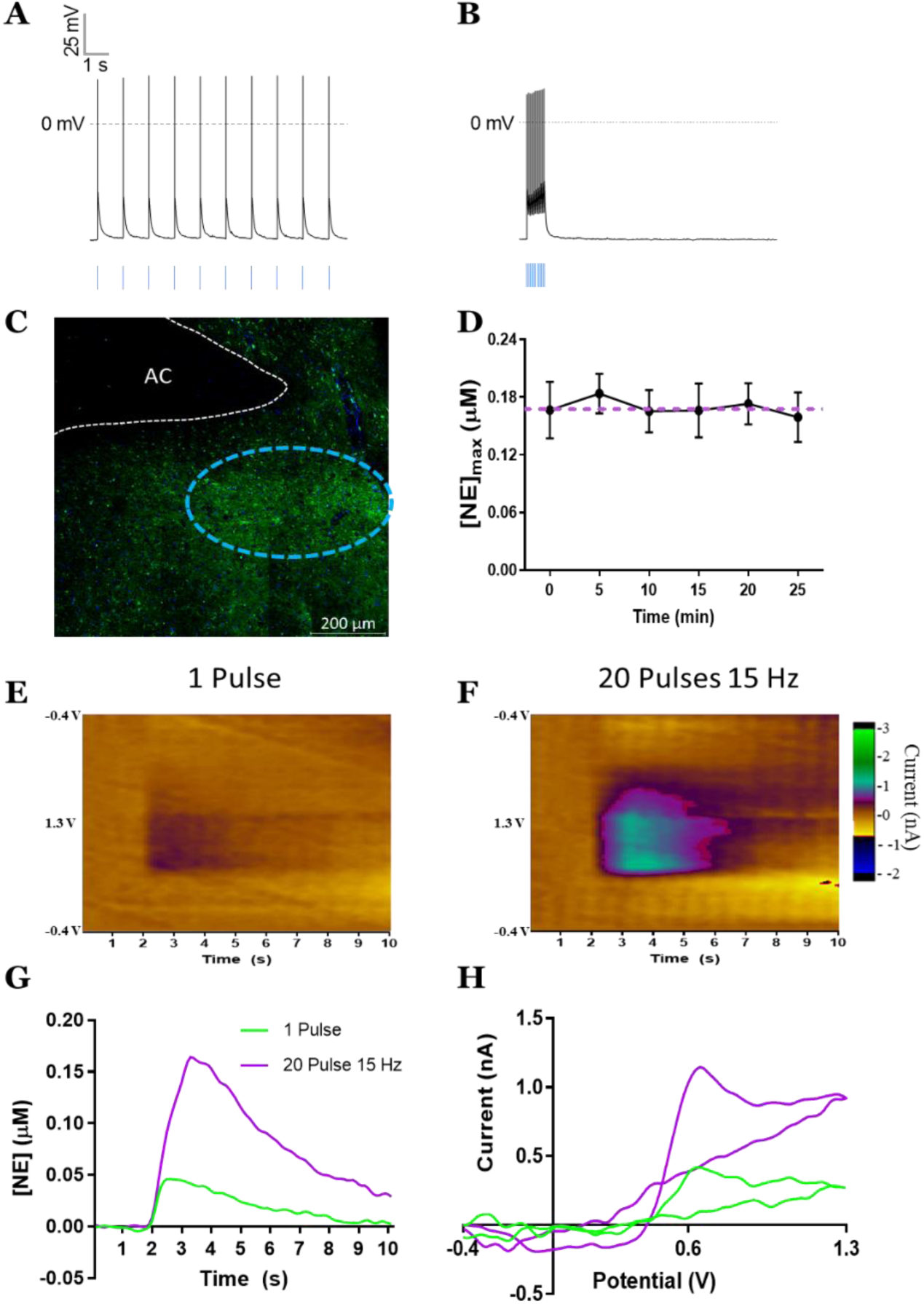
DBH:Cre(+/-)::Ai32(+/ +) mice act as a reliable model to study noradrenergic function. Patch clamp electrophysiology shows that GFP + neurons in A2 fire action potentials in response to optogenetic stimulation (blue bars) at both 1 Hz (A) and 15 Hz (B) frequencies. C) Terminal projections (green = GFP) in fusiform subnucleus (outlined in cyan dashes) of the ventral bed nucleus of the stria terminalis. D) 10 Hz, 20 pulse optogenetic stimulation of the vBNST induces consistent concentrations of norepinephrine release over multiple stimulations. Mean±SEM; n = 4 recording sites from 4 mice; p = 0.297. E) Color plot representing NE release following single pulse of optogenetic stimulation. F) Color plot representing NE release following 20 pulses at 15 Hz of optogenetic stimulation. G) Representative norepinephrine concentration over time following either single pulse (lime) or 20 pulses at 15 Hz (lilac). Optogenetic stimulation occurred at 2 s. H) Representative cyclic voltammograms of either single pulse (lime) or 20 pulses at 15 Hz (lilac).

After validating the utility of the DBH:Cre(+/-)::Ai32(+/ +) line to study the release and uptake of NE in the vBNST, we next examined if *in vivo* stress exposure would engage plasticity within this circuit. To ensure that our paradigm engaged norepinephrine neurons and elicited a stress response, we exposed mice on the same background to a 2-h restraint stress paradigm. Immediately following a restraint stress session, we collected trunk blood and analyzed it for plasma corticosterone (CORT) concentrations. As anticipated, restraint stress exposure significantly elevated CORT (Fig. 2A). Simultaneously, we retrieved the brains from these mice and used dual fluorophore fluorescent *in situ* hybridization (FISH) to probe for *cFos* and *DBH* mRNA in the nucleus of solitary tract. Restraint stress increased *cFos* translation in the A2 NE neurons (Fig. 2B). These data suggest that our stress paradigm stimulates activity in NE neurons and engages the hypothalamic-pituitary-adrenal stress axis.

**Figure 2.**
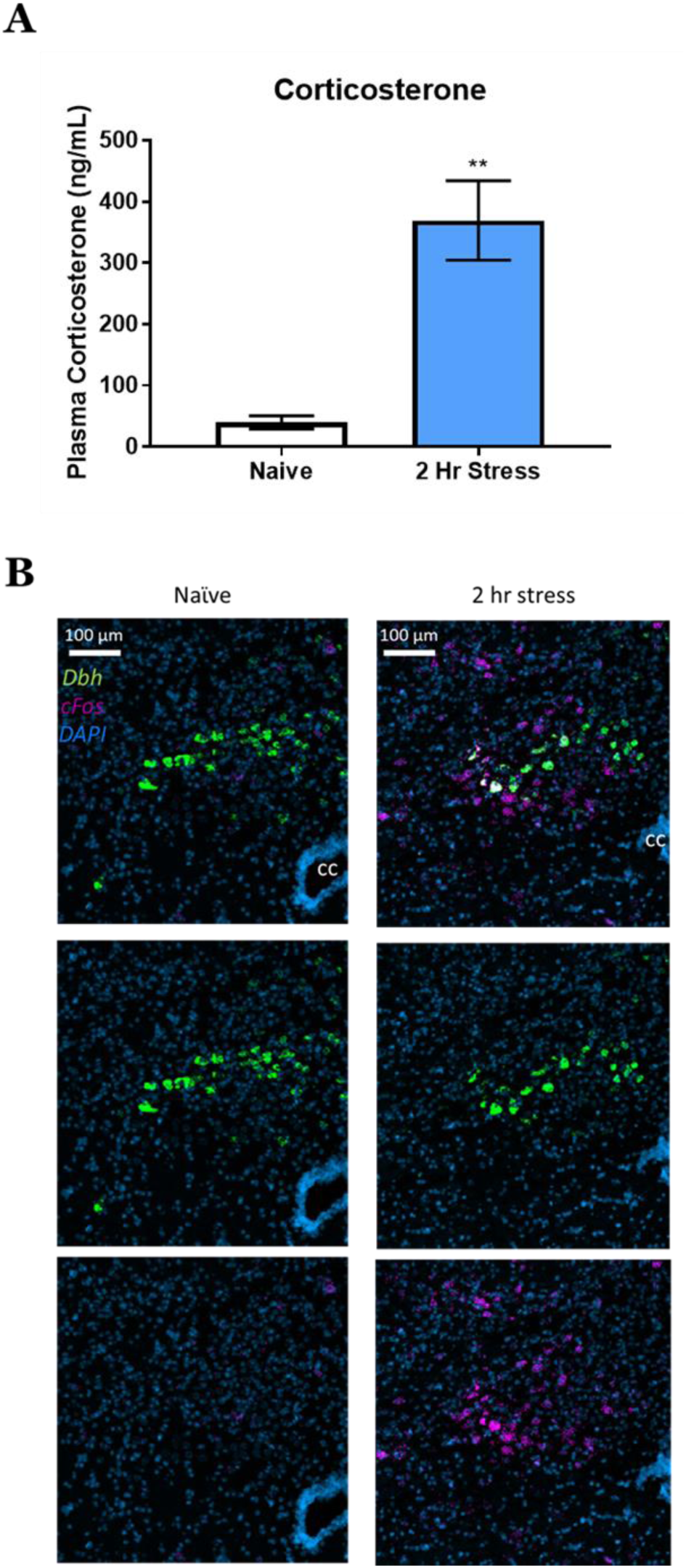
Two hours of restraint increases corticosterone and cFos expression. A) Plasma corticosterone (ng/mL) from naïve and 2 hr stress mice. Mean±SEM; n = 5 mice/group. Unpaired t-test, t = 5.027; p = 0.001. B) Nucleus of the solitary tract double fluorescent in situ hybridization showing DBH, cFos, and DAPI in both stress naïve and restraint stressed mice. CC = central canal.

We then performed slice optogenetics assisted FSCV to measure NE release in the BNST of mice from one of three conditions: stress naïve, single 2-hour restraint stress exposure, or repeated 5 days, 2-hour restraint stress exposure. All stress mice were recorded from on the day immediately following stress exposure to examine an adaptive response. Our stimulation parameters examined various frequency (20 pulses at 1, 2, 5, 10, 15 Hz) and pulse number (1, 2, 5, 10, 20 pulses at 5 Hz or 10 Hz) protocols, and were chosen to encapsulate a range of physiologically relevant noradrenergic firing patterns^24-26^. Repeated restraint stress exposure significantly increased NE release across a range of stimulation frequencies with the largest differences observed at high phasic frequencies (Fig. 3A). Stress naïve and single stress exposed animals did not significantly differ from one another. Similar results were observed with 5 Hz (Fig. 3B) and 10 Hz (Fig. 3C) phasic stimulations with the larger differences observed with increased pulses (approximately 3.3- to 5- fold difference following repeated stress exposure as compared to naïve and single stress conditions).

**Figure 3.**
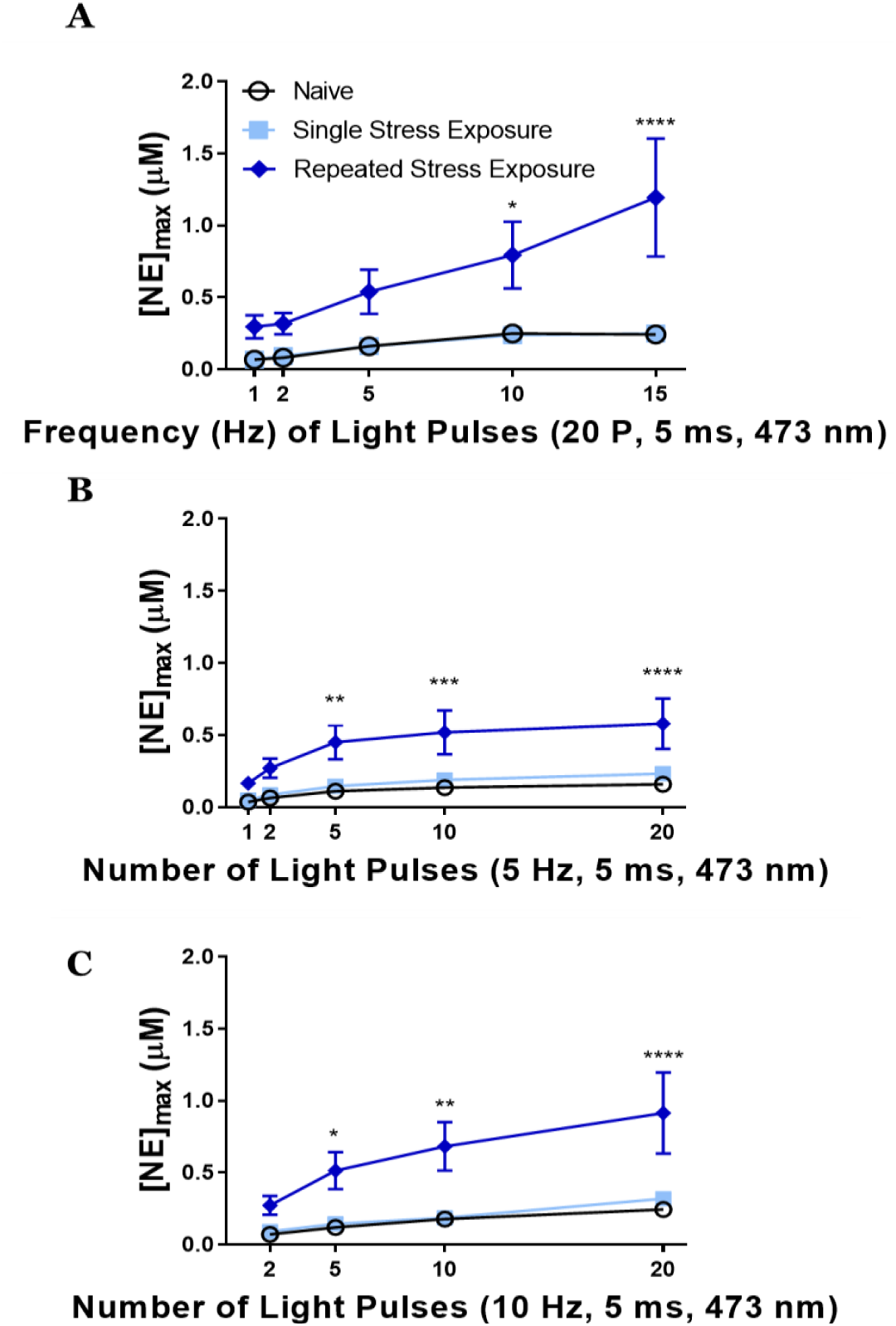
Repeated stress exposure increases norepinephrine release at multiple stimulation parameters.A) Varied frequencies of 20 pulse stimulations induce different peak concentrations of norepinephrine ([NE]max) between stress naïve, single stress exposed, and repeated stress exposed mice. Two-way repeated measures ANOVA, main effects of frequency (F = 7.66, p < 0.0001) and stress exposure (F = 7.733, p = 0.004) with a significant interaction (F = 2.484, p = 0.0199). B) Varied number of light pulses at 5 Hz stimulation induce different peak concentrations of norepinephrine between stress naïve, single stress exposed, and repeated stress exposed mice. Two-way repeated measures ANOVA, main effects of frequency (F = 23.01, p < 0.0001) and stress exposure (F = 7.37, p = 0.004) with a significant interaction (F = 3.229, p = 0.0031). C) Varied number of light pulses at 10 Hz stimulation induce different peak concentrations of norepinephrine between stress naïve, single stress exposed, and repeated stress exposed mice. Two-way repeated measures ANOVA, main effects of frequency (F = 21.27, p < 0.0001) and stress exposure (F = 8.09, p = 0.0027) with a significant interaction (F = 3.721, p = 0.0033). Mean±SEM; n = 7-8 slices/group from 5-6 mice/group. *p<0.05, **p<0.05, *** p<0.005, and ****p<0.0001.

Finally, in order to examine the role α_2_-adrenergic receptors may play at the noradrenergic terminals in the vBNST, we perfused the slices with the selective α_2_-adrenergic antagonist idazoxan. In both naïve and single stress exposed mice, 10 μM idazoxan significantly increased NE release (Fig. 4A and 4B) across a range of stimulation frequencies. Following repeated restraint stress, however, α_2_-antagonism failed to alter NE release (Fig. 4C). These data suggest that a loss in sensitivity at the α_2_-autoreceptor may underlie increased release of NE in mice that have experienced chronic stress. Furthermore, these data are consistent with experiments conducted in rat examining how morphine withdrawal and social isolation stress regulate NE release^17-18^, suggesting that this may be a common mechanism to elevate NE levels across species following traumatic stress.

**Figure 4.**
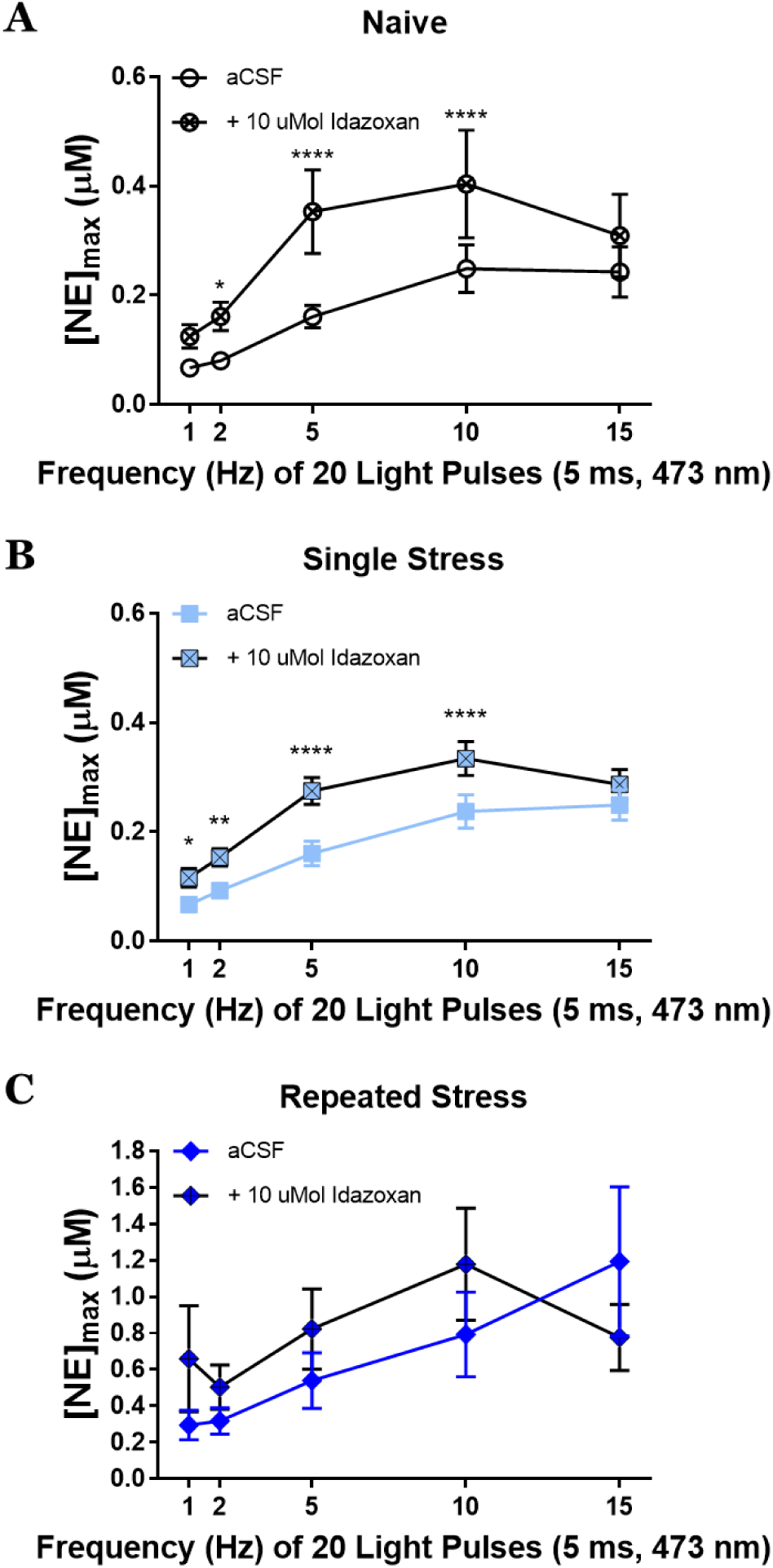
The α2-adrenergic autoreceptor antagonist, idazoxan (10 μMol), increases peak concentrations of norepinephrine ([NE]max) in stress naïve (A), F= 11.05, p = 0.02, and single stress (B), F = 74.18, p = 0.0001, but not repeated stress (C) mice, p > 0.05. aCSF = artificial cerebrospinal fluid. Mean±SEM; n = 6-7 slices/group from 5-6 mice/group. *p<0.05,**p<0.05, *** p<0.005, and ****p<0.0001.

The alterations of noradrenergic release properties we have shown may illustrate important underlying neurobiological mechanisms of psychiatric disorders. We all experience stress to some degree in our daily lives, but the transition from “normal” stress to maladaptive stress likely alters neuronal systems. Our data here, indicate that one of these systems altered by repeated stress exposure is the noradrenergic system which shows greater response to optogenetic stimuli and less control by inhibitory autoreceptors. A greater understanding of how this transition occurs, such as the duration or intensities of stress exposures necessary to alter NE function, the persistence of this effect, and additional mechanisms that contribute to these adaptations, are still needed to further enhance our knowledge of stress disorders.

Furthermore, our data here show that the DBH:Cre(+/-)::Ai32(+/ +) mouse will be a useful tool to continue to investigate these processes, both in terms of adaptation within the noradrenergic circuitry itself, and in the subsequent neural systems that are modulated by NE. Regardless, this elevated noradrenergic tone may relate to hyperarousal states in disorders such as PTSD^4^, altered affect^1^, and susceptibility for addiction^3^.

## Methods

### Animals and Housing

Adult male and female mice were tested with sex counterbalanced across groups. Selective expression of ChR2 in noradrenergic neurons was achieved by breeding a cross between DBH:Cre(+/-)::Ai32(+/ +) mouse with a Ai32(+/ +) mouse. Mice were housed with littermates in a temperature controlled vivarium with a 12:12 h light-dark cycle with lights on at 0700 h, and had *ad libitum* access to food and water. All procedures were approved by the Institutional Animal Care and Use Committee of the University of North Carolina at Chapel Hill.

### Stress Procedures

Mice were placed in custom in-house modified ventilated 50 mL conical tubes for 2 h. Mice in the single stress exposure condition were tested the day following restraint. Mice in the repeated stress exposure condition were underwent restraint procedures for 2 h/day for 5 consecutive days. These mice were tested the day following the fifth restraint session.

### Corticosterone Analysis

Mice were decapitated following restraint stress or from homecage (at 12:00 pm) and trunk blood was collected into heparinized tubes. Blood samples were immediately centrifuged (2000 × g for 10 minutes), plasma was isolated, placed on dry ice, and stored at −80°C until analysis. Plasma was analyzed for corticosterone content using a commercially available colorimetric ELISA kit (Arbor Assays; Ann Arbor, MI), according to the manufacturer’s instructions. All samples were run in duplicate.

### Double Fluorescent *In Situ* Hybridization (FISH)

Brains were rapidly removed and flash frozen on dry ice for a minimum of 5 minutes, and then stored at −80 °C for no more than 1 week prior to slicing. Sections were then sliced on a cryostat (Leica 300s, Germany) at 18 μm and directly mounted onto slides, and stored at −80 °C prior to the FISH procedure. FISH was performed using the RNAScope kit (ACD Biotechne) according to the manufacturer’s instructions (except the time for protease IV step was reduced to 15 min) using antisense probes against *Dbh* (Mm-Dbh-C2 400921-C2) and *cFos* (Mm-Fos-C3 316921-C3). Sections were imaged on a Zeiss 800 confocal microscope using identical settings.

### Slice Preparation, Electrophysiology, and Electrochemistry Recordings

Mice were deeply anesthetized (isoflurane), decapitated, and brains were harvested and placed in ice-cold sucrose for slicing (ACSF in mM: 194 sucrose, 20 NaCl, 4.4 KCl, 2 CaCl_2_, 1 MgCl_2_, 1.2 NaH_2_PO_4_, 10 glucose, 26 NaHCO_3_) that had been oxygenated with 95% O2, 5% CO2 for at least 15 min. Brains were sliced at 300 μm using a Leica VT1000 (Germany). Following slicing, brains were incubated in oxygenated ACSF (in mM: 124 NaCl, 4.4 KCl, 2 CaCl_2_, 1.2 MgSO_4_, 1 NaH_2_PO_4_, 10 glucose, 26 NaHCO_3_, 34§ C) and allowed to incubate for at least 30 minutes. Slices were transferred to the electrophysiology or electrochemistry rigs for patch clamp or fast scan cyclic voltammetry, respectively. Each rig perfused oxygenated ACSF (28-30 °C) through the bath at 2 ml/min. Cells expressing ChR2-eYFP were acquired using whole-cell voltage clamp and then switched to current-clamp mode (in mM: 135 gluconic acid-potassium, 5 NaCl, 2 MgCl_2_, 10 HEPES, 0.6 EGTA, 4 Na_2_ATP, 0.4 NA2GTP). Because NE neurons are spontaneously active, cells current was injected to keep the membrane potential at −70 mV and 10, 5 ms light pulses were applied with varying frequency (1, 2, 5, 10, and 15 Hz.) Electrochemical recordings were made as described previously^27^. Carbon fiber microelectrodes were fabricated in house with fiber lengths 50-100 μm. Electrodes were calibrated with 3 concentrations of NE: 0.1 μM, 1.0 μM, and 10 μM NE. By graphing the peak current induced by each concentration of NE, we could draw a line of best fit whose slope was used as the current-concentration calibration factor for that electrode. Using a custom built potentiostat (University of Washington, Seattle) and TarHeel CV written in laboratory view (National Instruments), a triangular waveform (−0.4 V to 1.3 V) was applied at 10 Hz. Slices were optically stimulated with 5-ms blue (490 nm) light pulses down the submerged 40x objective. Stimulation parameters included single pulse; 2, 5, 10, 20 pulses at either 5 Hz or 10 Hz; and 20 pulses at 1, 2, 5, 10, and 15 Hz. Each recording began with 2 s (20 voltammograms) of recording before stimulus delivery. Recordings were separated by >5 min. Voltammograms were analyzed with HDCV (UNC Chapel Hill) with NE currents isolated using principal component regression analysis, as described previously^28^, and [NE]_max_ was determined with Clampfit 10.6 software (Molecular Devices, Sunnyvale, CA). 10 μM idazoxan (Sigma-Adrich, St Louis, MO) in ACSF was bath applied for >10 min before testing.

### Data Analysis and Statistics

Data were analyzed using GraphPad Prism 7.0 and are represented as mean ± SEM. CORT comparisons were conducted using an unpaired Student’s *t-*test. Two-way ANOVAs were performed on all NE release analyses with repeated factors of Frequency (Fig. 3A and 4A) or Number of Light Pulses (Fig. 3B and C; 4B and C) and Drug Treatment (Fig. 4). Tukey’s or Sidak’s post hoc tests were conducted where appropriate. Significant differences were represented at \**p<*0.05, \*\**p<*0.05, *** *p<*0.005, and \*\*\*\**p<*0.0001.

## Supporting Information

### Abbreviations

ACSF: artificial cerebrospinal fluid
BNST: bed nucleus of the stria terminalis
ChR2: channelrhodopsin
CORT: corticosterone
FSCV: fast-scan cyclic voltammetry
NE: norepinephrine.

## Author Information

Tel: (919)966-8637

## Author Contributions

Concept and design, Z.A.M. and K.T.S.; K.T.S. responsible for electrode fabrication, FSCV experiments, data analysis, writing the manuscript; V.H.M and J.B. responsible for CORT analysis; K.M.B. responsible for FISH; D.P. and M.M.P. responsible for electrophysiological recordings; I.M.B participated in stress procedures, electrode fabrication, and FSCV experiments; J.L.L. and S.R.J responsible for HPLC; Z.A.M. responsible for writing the manuscript.

## Acknowledgements

The authors would like to thank Dr. Patricia Jensen for generating and providing the DBH::Cre mice for these experiments. We also thank the Hooker Imaging Core at UNC and support from the Bowles Center for Alcohol Studies at UNC. We would like to thank Dr. Thomas Kash for comments on a previous version of the manuscript.

